# A microbial metabolite reduces alcohol-induced inflammation via dual modulation of NF-κB and Interferon pathway

**DOI:** 10.64898/2026.06.18.733199

**Authors:** Yurui Zheng, Nicholas Handali, Dana Moradi, Colin Varnet, Farhin Patel, Alexander Aksenov, Adam Kim

## Abstract

**Background and aims:** Alcohol-associated hepatitis (AH) is characterized by excessive inflammation and blunted antiviral interferon (IFN) responses. We hypothesized that specific gut microbiome-derived metabolites could selectively enhance interferon signaling while limiting NF-κB mediated inflammation, thereby restoring immune balance in AH. Our goal is to identify microbiome-derived metabolites that differentially regulate the NF-κB and IFN signaling pathways.

**Methods and results:** We used human monocytic THP1-Dual cells, which secrete reporters for NF-κB and IFN signaling, to model innate immune responses and screened a library of 152 gut microbiome-derived metabolites. From the metabolite screen, 4-hydroxyphenylacetic acid (4-HPAA) emerged as a unique immunomodulator: in LPS-challenged cells, 4-HPAA selectively increased IFN signaling with minimal NF-κB activation. 4-HPAA was evaluated *in vivo* using a NIAAA-model, with 4-HPAA supplementation (0.4mg/ml) added to the diet. In the NIAAA-model, dietary 4-HPAA did not induce liver injury and was associated with enhanced interferon-stimulated gene expression. Simultaneously, 4-HPAA reduced pro-inflammatory markers such as *Il1β*, *Ly6g* and *F4/80* compared to the group exposed to ethanol alone. Metabolomic profiling of mouse cecal contents revealed 4-HPAA supplementation counteracted ethanol’s metabolic effects, selectively reducing triglyceride-associated lipids that had accumulated with ethanol feeding.

**Conclusions:** 4-HPAA enhances interferon signaling and antiviral gene induction while dampening NF-κB-driven inflammation in the presence of LPS, both *in vitro* and *in vivo*. In an acute-on-chronic alcohol injury model, 4-HPAA attenuated hepatic inflammation, reduced immune cell recruitment, and activated antioxidant defenses, reflecting a shift toward a more hepatoprotective effect. 4-HPAA treatment was associated with reduced pro-inflammatory markers and modest attenuation of ethanol-induced liver injury. Additionally, 4-HPAA reversed ethanol-induced lipid-dysregulation, particularly triglyceride accumulation, highlighting its metabolic benefit in alcohol-fed mice. In conclusion, 4-HPAA rebalances immune and metabolic pathways by enhancing IFN signaling, suppressing NF-κB inflammation, and reversing alcohol-induced hepatic injury and lipid accumulation.

## Introduction

Chronic alcohol consumption is a major factor in the development of alcohol-associated liver disease (ALD), which ranges from steatohepatitis, fibrosis, cirrhosis, to hepatocellular carcinoma[1]. Ethanol metabolism in hepatocytes generates acetaldehyde and reactive oxygen species[2], leading to oxidative stress, lipid peroxidation[3], and mitochondrial dysfunction[4]. Concurrently, alcohol disrupts intestinal barrier integrity and alters gut microbiota composition[5], promoting translocation of bacterial products such as lipopolysaccharide (LPS) into portal circulation[6]. These microbial components activate Kupffer cells via Toll-like receptor 4 (TLR4), triggering NF-κB signaling and the release of pro-inflammatory mediators including tumor necrosis factor alpha (*TNF-α*) and interleukin 6 (*IL-6*). TLR4 activation also triggers a TRIF-dependent pathway that induces interferon (IFN) signaling via IRF3 and IRF7[7]. Alcohol is well known to alter NF-κB signaling, contributing to hepatic inflammation and immune dysregulation independently[8]. However, its impact on IFN signaling is less well understood, and recent evidence suggests that ethanol may also disrupt this pathway, further impairing antiviral immune responses in alcohol-associated hepatitis (AH) patients[9, 10]. Sustained immune activation and hepatocellular injury promote stellate cell activation and extracellular matrix deposition, driving fibrosis and disease progression[11]. Thus, chronic alcohol intake establishes a pro-inflammatory hepatic environment through metabolic stress, gut-derived endotoxin signaling, and immune dysregulation, forming the mechanistic basis of ALD pathogenesis[12].

Microbiome-derived metabolites play a critical role in modulating immune responses during chronic alcohol consumption and the progression of ALD[13]. Dysbiosis induced by ethanol consumption alters microbial metabolic activity[14] which in turn affects production of short-chain fatty acids (SCFAs) and bile acids[15]. SCFAs, in particular butyrate, normally maintain gut barrier integrity and exert anti-inflammatory effects through G-protein–coupled receptor such as GPR43 signaling [16] and inhibition of histone deacetylases[17]. Similarly, alterations in bile acid profiles disrupt FXR and TGR5 signaling pathways, which regulate bile acid synthesis, lipid metabolism, and inflammatory responses in the liver[18].

In this study, we sought to identify microbiome-derived metabolites that differentially regulate the NF-κB and IFN signaling pathways. We hypothesized that the microbiome would exert protective efects against external stressors such as alcohol consumption, whereby specific microbiome-derived metabolites can selectively enhance IFN signaling while limiting NF-κB activation, thereby restoring immune balance. To test this, we screened a defined microbial metabolite library in THP1-Dual cells and identified 4-hydroxyphenylacetic acid (4-HPAA), a gut microbial metabolite derived from dietary tyrosine, as a candidate immunomodulator for further *in vivo* investigation.

## Methods

### THP1-Dual cell culture

THP1-Dual™ (InvivoGen, USA) cell line was cultured in RPMI 1640 medium containing 2 mM L-glutamine, 25 mM HEPES, 10% heat-inactivated fetal bovine serum (30 min at 56°C), 100 μg/ml Normocin and 100 U/ml Pen-Strep. To maintain selection pressure, 10 μg/mL of Blasticidin and 100 mg/mL of Zeocin antibiotics were added to the growth medium every other passage.

### Ethanol test

THP1-Dual™ cell was plated in 96-well plate at concentration of 5.56×10^5^ cells/ml per well. Cells (n=3) were challenged with ethanol (0, 10mM, 20mM, 50mM, 100mM). After 24 hours, 100pg/ml LPS was added to cells. After 24 hours incubation, supernatant was collected at 300 x g for 5mins. SEAP and luciferase results were collected from cell supernatant by following manufacturer instructions.

### Metabolite screening test

THP1-Dual™ cells were plated in a 96-well plate at a concentration of 5.56×10^5^ cells/ml (n=3 for all experimental conditions). Cells were challenged with +/- LPS (100pg/ml). After 24 hours incubation, cells were treated with one compound from a microbiome metabolite library, 152 compounds in total (Cayman Chemical, USA). After 24 hours incubation, cells were pelleted at 300 x g for 5mins, then supernatant was collected. SEAP and luciferase results were measured.

### Mice

All animals received humane care. WT C57BL/6J mice were purchased from The Jackson Laboratory (Bar Harbor, ME, USA). Mice were allowed free access to a Lieber-DeCarli liquid diet containing +/- ethanol at 5% (v/v) and +/- 0.4 mg/ml 4-HPAA that isocalorically substituted maltose dextrin for ethanol for 10 days. On the final day of the experiment, pair-fed mice were gavaged with 5g/kg maltose and ethanol-fed mice were gavaged with 5g/kg ethanol in water. Mice were sacrificed 6 hours after the ethanol binge, a commonly used time point in the NIAAA model to capture peak acute liver injury and inflammatory responses[19].

### Isolation of bone marrow macrophage from mice

3-month-old wild-type mice (4 males and 4 females)were euthanized under approved protocol. Femurs and tibias were dissected from mice and cleaned in sterile PBS. One end of each bone was cut off. 0.5ml tube cap was cut away and a small hole was made by an 18G syringe needle in the bottom of 0.5ml tube. Each bone was put in a 0.5ml centrifuge tube with the side that was cut facing down and entire tube was transferred into a 1.5ml centrifuge tube. The 1.5ml tube was centrifuged at 10,000 rpm for 1 second. 1.5ml tube containing bone marrow was kept. 1 ml bone marrow culture media was added to the 1.5ml tube and suspended the cells. Then all cells were pipetted into a 50ml tube which had 40ml of media through a 35μm filter. Distribute 1ml of cells into 24-well plate. For macrophage differentiation, cells were cultured in RPMI 1640 supplemented with M-CSF (20 ng/mL) at 37 °C with 5% CO₂ for 10 days, and the medium was replaced every 3 days. On day 11, cells were treated with +/- 10μ M 4-HPAA. After 24 hours, cells were centrifuged at 300 x g for 5mins to remove supernatant.

200μl of Trizol was added to cells and cells were stored in -20℃ for further RNA isolation. Mouse of origin was treated as biological replicate, and treatment comparisons were paired by donor. Sex was included as a biological variable in the experimental design.

### Biochemical assays

Plasma samples were assayed for alanine aminotransferase (ALT) and aspartate aminotransferase (AST) using a commercially available enzymatic assay kit (Sekisui Diagnostics, Lexington, MA), following the manufacturer’s instructions.

### Histopathology

Formalin-fixed liver tissues were paraffin-embedded, sectioned and stained with hematoxylin and eosin for histological analysis. Tissues were coded at time of collection to assure an unbiased analysis; 2 images were acquired per tissue section.

### RNA extraction and Quantitative Real-Time Polymerase Chain Reaction(qRT-PCR)

RNA was extracted by using the Direct-zol mini kit per the manufacturer’s instructions (Zymo Research, Irvine, CA). For mRNA measurement, RNA was reverse-transcribed using the Superscript Vilo kit (Thermo Fischer, Waltham, MA) and measured with PowerSYBR qRT-PCR kits (Applied Biosystems, Foster City, CA) on a QuantStudio5 analyzer (Applied Biosystems, Foster City, CA). Relative messenger RNA (mRNA) expression was determined using gene-specific primers. Analyses were performed using the ΔΔCt method.

### Bulk RNA sequencing of BMDMs

We extracted RNA from BMDMs using the Zymo Direct-zol kit with an in-column DNAse step. 1 ug of RNA was used to prepare Illumina stranded RNA-seq libraries using the SMART-Seq® Total RNA High Input (RiboGone™Mammalian) kit (Takara Bio, San Jose, CA). Libraries were multiplexed and sequenced on Illumina NovaSeq to generate more than 2 billion paired-end reads of 100 bp with a median of 18 million reads per sample.

### Analysis of bulk RNA sequencing from patients’ liver

RNA-seq data was obtained from dbGAP (phs001807.v1.p1)[20]. Raw fastqs were aligned to the human genome (GRCh38, release 96) and gene expression was determined using kallisto in gene mode with 100 bootstraps[21]. Differential gene expression was measured using Sleuth in gene mode using the likelihood ratio test (LRT) with a cutoff of FDR<0.05[22]. For all figures and deconvolution, bootstrap data were converted to TPMs in Sleuth. Heatmaps were generated in Sleuth, scaled by row and sample clustering turned off.

### Analysis of bulk RNA sequencing from BMDMs

Fastq files were aligned to the mouse genome (GRCm39, release 113) using Kallisto version 0.46.1 with 100 bootstraps calculated[21]. Data were then merged with experimental data and analyzed with Sleuth in gene_mode with aggregation_column set to Ensemble Gene ID; in addition, extra_bootstrap_summary and read_bootstrap_tpm were set to true[22]. Differential expression was measured with Sleuth using a cutoff of q<0.05.

### Analysis of bulk RNA-sequencing data from mouse liver

Raw sequencing data (fastq files) were obtained from the original publication (GSE188967)[23]. Fastq files were aligned to the mouse genome (GRCm39, release 113) using Kallisto version 0.46.1 with 100 bootstraps calculated[21]. Data were then merged with experimental data and analyzed with Sleuth in gene_mode with aggregation_column set to Ensemble Gene ID; in addition, extra_bootstrap_summary and read_bootstrap_tpm were set to true[22]. Differential expression was measured with Sleuth using a cutoff of q<0.05.

### Metabolomics analysis

The samples were thawed at room temperature for 10 minutes. Each empty Eppendorf tube was placed on the balance, the balanced tared, and the sample transferred into the tube. The weight of the sample was recorded. An aliquot of HPLC-grade 80% v:v ethanol: water at a ratio of 10 µL per 1 mg of sample was added into each tube. The material was homogenized at 90Hz for 5 min using Precellys® 24 Touch tissue homogenizer (Lysis & homogenization). The samples were allowed to be extracted overnight at room temperature. The Eppendorf tubes were Centrifuged for 10 minutes at 2,000 and 100μl of supernatant from each tube transferred to a fresh Eppendorf and dried down using Integrated Speedvac System Iss110. Then, re-suspended in 100μl of 50% v/v ethanol, sonicated for 10min, and centrifuged at 2,000r.p.m. for 10min. A 30 μl aliquot of each sample was transferred to a vial, and 150 µl of HPLC-grade 90% v:v acetonitrile was added. Also, blanks of resuspension solvent as well as the solvent that went through the extraction procedure steps (sample extraction blank) were included with each batch.

### LC-MS/MS data acquisition and processing

Blank solvent injections were carried out every 10 samples. Sample extraction blanks and QC of four sulfa drugs were included. Vanquish UPLC (Thermo Fisher Scientific, Waltham, MA), was used for injecting samples and chromatographic separation using C18 chromatography column, 50mm × 2.1 mm Kinetex 1.7 μM, C18, 100 Å (Phenomenex, Torrance, CA), 40 °C column temperature, 0.4 mL/min flow rate, mobile phase A 99.9% water (Fisher Scientific, Optima LC–MS), 0.1% formic acid (Thermo Fisher Scientific, Optima LC/MS), mobile phase B 99.9% acetonitrile (Fisher Scientific, Optima LC–MS), 0.1% formic acid (Fisher Scientific, Optima LC–MS). The solvent gradient table was set as follows: 0.2% B, increased to 10% B over 0.2min, then to 100% B at 12min, held at 100% B for 1.5min, decreased back to 2% B in 0.5min, followed by washout cycle and equilibration for a total analysis time of 14min.

MS analysis in positive polarity mode was performed on Orbitrap Exploris 480 (Thermo Fisher Scientific, Waltham, MA) mass spectrometer equipped with H-ESI-II probe sources. The following probe settings were used for both MS for flow aspiration and ionization: spray voltage of 3500 V, sheath gas (N2) pressure of 35 psi, auxiliary gas pressure (N2) of 10 psi, ion source temperature of 350 °C, S-lens RF level of 50 Hz and auxiliary gas heater temperature at 400 °C.

Spectra were acquired in positive ion mode over a mass range of 75–1125 Th. An external calibration with Pierce LTQ Velos ESI Positive ion calibration solution (Thermo Fisher Scientific, Waltham, MA) was performed prior to data acquisition with ppm error of less than 1. Data was recorded with data-dependent MS/MS acquisition mode. Full scan at MS1 level was performed with 30K resolution. MS2 scans were performed at 11250 resolutions with max IT time of 60 ms in profile mode. MS/MS precursor selection windows were set to m/z 1.5 with m/z 0.5 offset. MS/MS active exclusion parameter was set to 5.0 s.

### Metabolomics data processing

All raw data are publicly available at massive.ucsd.edu; dataset ID MSV000098166. The feature tables were obtained using MZmine4[24]. The collected HPLC-MS raw data was converted from Thermo’s .raw to .mzML format. The data were then batch-processed with the following settings for each step: mass detection (noise level 1000), chromatogram builder (minimum time span 0.01 min, minimum peak height 3000, m/z tolerance 0.1 m/z or 20 ppm), chromatogram deconvolution - baseline cutoff (minimum peak height 3000, peak duration range 0.01-3.00 min, baseline level 300), deisotopisation - isotopic peak grouper (m/z tolerance 0.1 m/z or 20 ppm, RT tolerance 0.1 min, maximum charge 4), peak alignment - join aligner (m/z tolerance 0.1 m/z or 20 ppm, weight for m/z 75, weight for RT 25, RT tolerance 0.1 min), peak filtering - peak list raw filter (minimum peak in a row 3, minimum peak in an isotope pattern 2).

### Mass spectrometry statistical analysis

MetaboAnalyst [25] was utilized for Principal Component Analysis (PCA), Partial Least Squares-Discriminant Analysis (PLS-DA) and Cross Validation to evaluate the variability of metabolites introduced by different diets. Features with values below 3x of the corresponding feature in blanks were removed (blank subtraction). Zero values in the feature table were imputed by replacing them with one-fifth minimum positive value for each feature. The parameters for Data Filtering were set as: 25% for Reliability filter, Interquartile range (IQR) for Variance filter and mean intensity value for Abundance filter. Also, data normalization parameters were set as Quantile normalization for Sample normalization, Log transformation for Data transformation and Auto scaling for Data scaling.

### Molecular Networking

Cytoscape[26] was used to visualize a molecular community network (MCN)[27], which was constructed to organize and interpret the relational structure among detected metabolites from untargeted LC-MS/MS data. The feature table and MS/MS spectral file (MGF), generated by MZmine4, were submitted to the Feature-Based Molecular Networking (FBMN) workflow on GNPS platform (gnps.ucsd.edu)[28]. The data were clustered with MS-Cluster with a parent mass tolerance of 0.02 Da and an MS/MS fragment ion tolerance of 0.02 Da to create consensus spectra, with those containing fewer than three spectra discarded. A network was then created where edges were filtered to have a cosine score above 0.7 and more than 6 matched peaks. The spectra in the network were then searched against GNPS’s spectral libraries. Subsequently, a new job was cloned from the original GNPS job with parameters set as follows: minimum pairs cosine = 0.1, Network TopK = 100, and Maximum Connected Component Size = 0. Upon completion, the unpruned network (GraphML) was downloaded from the GNPS results page Molecular communities were then defined using the Louvain algorithm implemented via MCN scripts and the resulting community network was visualized and annotated in Cytoscape. The GNPS Jobs and the parameters used are available at the links below:

Original GNPS Job: https://gnps.ucsd.edu/ProteoSAFe/status.jsp?task=e0304fd3a3474379ad85fb9057148324

Unpruned Job used for MCN construction: https://gnps.ucsd.edu/ProteoSAFe/status.jsp?task=3edfe9029fa149d39561cf75fa3ea5fc

deposited code for Unpruned network: https://github.com/Alexander0/molecular_communities

### Statistical analysis

All statistical analyses were performed in R (v4.5.2). Data visualization was conducted using the ggplot2 package (v4.0.0). Normality of residuals was assessed using the Shapiro–Wilk test. All tests were two-tailed, and a threshold of *p* < 0.05 was used to determine statistical significance.

## Results

### ISGs are dysregulated in livers of AH patients

Interferon stimulated genes (ISGs) are an important aspect of innate immunity, most well known for their role in anti-viral defense. In viral infections like Hepatitis C, interferons and ISGs help eliminate the virus and damaged cells, but alcohol consumption can severely limit the effectiveness of interferon therapy in HCV patients[29], though interferon therapy is no longer standard of care[30]. Using bulk RNA-seq data from patients with different chronic liver diseases, we analyzed ISG expression to better understand how interferon signaling might be disrupted in AH[20]. These data include patients with early AH (EAH), severe AH with liver failure (AHL), severe AH from explant tissue (ExAH), non-alcoholic fatty liver disease (NAFLD), Hepatitis C virus (HCV), and HCV with cirrhosis (HCV-Cirr). As expected, many ISGs were upregulated in HCV and HCV-Cirr (**Figure 1**). Of the ISGs upregulated in HCV, some were significantly downregulated in AH (Ifit1) while others were also significantly upregulated but a much lower magnitude (*IFITM1*). Interestingly, other ISGs were only upregulated in AH (*IFITM2*). We hypothesized that this might be due to differential expresson of key negative regulators of interferon signaling, specifically suppressor of cytokine signaling (SOCS1), which is upregulated in AH, and Ubiquitin specific peptidase 18 (*USP18*), which is downregulated. *SOCS1* and *USP18* are both upregulated by type 1 and type 3 interferons as feedback inhibitors but have different mechanisms to negatively regulate signaling[31]. *SOCS1* inhibits JAK/Stat signaling, essential for ISG expression, and high *SOCS1* levels reduces interferon responses in tissues[32]. *USP18* only inhibits type 1 interferon responses, so other signaling can persist[31]. These data suggest that in AH, liver tissue is unable to express many ISGs, possibly due to higher *SOCS1* expression and reduced Jak/Stat signaling, leading to altered interferon signaling.

**Figure 1:**
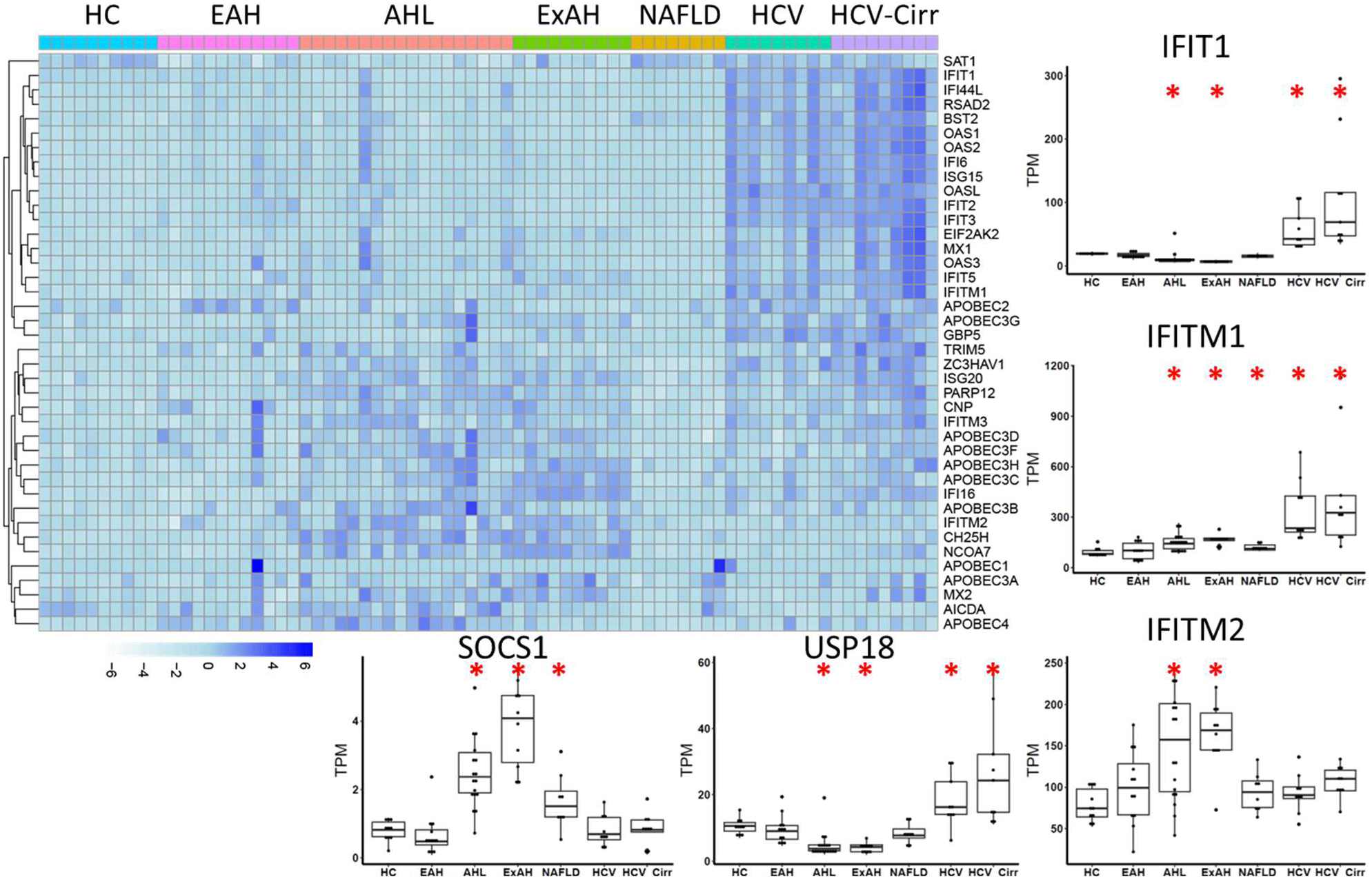
AH patients have different ISG expression compared to HCV. Bulk RNA-seq of liver isolated from patients with different chronic diseases. Left – Heatmap of ISG expression. Right - Boxplots of three ISGs with variable expression patterns in disease. Bottom - Boxplots of two genes which regulate ISG expression. HC-Healthy Control, EAH-Early AH, AHL-Severe AH with Liver Failure, ExAH-Explant tissue from severe AH, NAFLD-Nonalcoholic Fatty Liver Disease, HCVHepatitis C Virus, HCV-Cirr-HCV with Cirrhosis. * Indicates p<0.05 compared to HC.

### Low dose LPS and ethanol can induce imbalance in both NF-κB pathway and Interferon signaling pathway

Previous studies showed that monocytes in AH respond to low-dose LPS (100pg/ml LPS) with increased NF-κB signaling and reduced IFN signaling[9]. To confirm combined effects of both low dose LPS (100pg/ml LPS) and ethanol, we challenged THP1-Dual cells with ethanol at different concentrations (0, 10mM, 20mM, 50mM, 100mM) on day 1. After 24 hours, we treated cells with low dose LPS (+/- 100 pg/ml), which is comparable to endotoxin levels after binge alcohol consumption and in patients with AH[33, 34]. THP1-Dual cells are a human monocytic cell line engineered to simultaneously monitor activation of the NF-κB and type I interferon (IFN-I) signaling pathways. LPS signaling through TLR4 leads to NF-κB activation and transcription of proinflammatory genes and a secreted embryonic alkaline phosphatase (SEAP) reporter gene, which enables quantification of NF-κB activity in the culture supernatant. In parallel, endosomal TLR4 signaling induces transcription of interferon-stimulated genes and secreted luciferase gene under the control of an interferon-stimulated response element (ISRE). As expected, low-dose LPS (100pg/ml LPS) lead to both SEAP and luciferase secretion, indicating activation of both pathways. Ethanol was tested across a broad concentration range (0–100 mM) to assess dose responsiveness. Among these concentrations, 10–20 mM is more physiologically relevant to moderate alcohol exposure[35], whereas 50–100 mM represents higher experimental exposure and should be interpreted primarily as mechanistic in vitro conditions. Ethanol pre-treatment enhanced NF-κB activation in a dose-dependent manner (**Figure 2**). In contrast, pre-treatment with ethanol resulted in a progressive decrease in luciferase activity, suggesting that ethanol suppressed interferon signaling in a dose-dependent manner. These results indicate that ethanol skewed LPS-induced immune responses by amplifying NF-κB–driven inflammation while dampening IFN signaling, potentially contributing to an imbalanced immune state.

**Figure 2:**
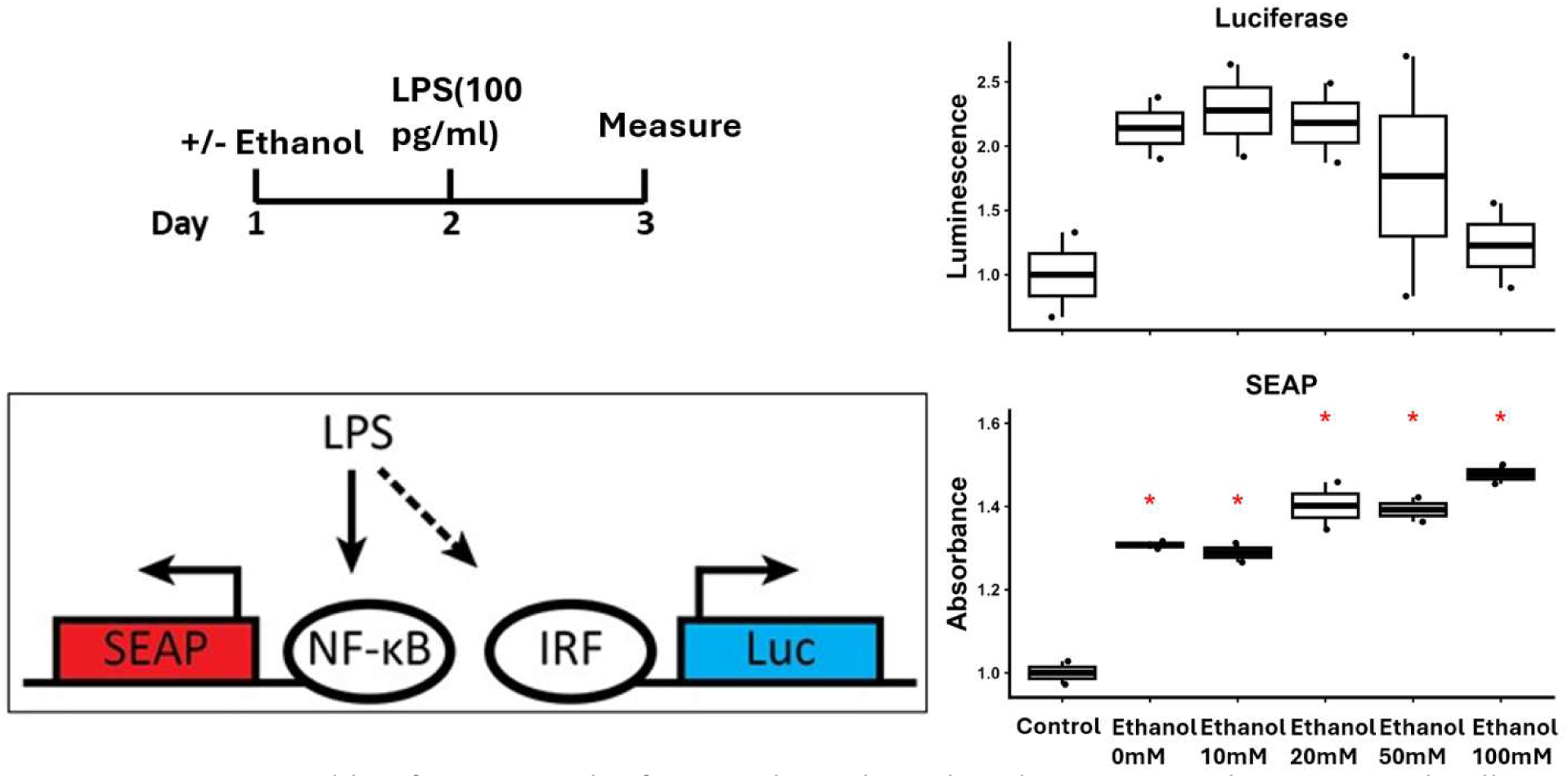
SEAP and luciferase results from 3-day ethanol and LPS test with THP1-Dual cells. 5 different ethanol concentrations were tested with 100pg/ml LPS: 0, 10 mM, 20 mM, 50 mM, 100 mM. Day 1, THP1-Dual cells were treated with or without ethanol; Day 2, THP1- Dual cells were challenged with or without low dose LPS (100pg/ml LPS:); Day 3, cell supernatant was collected for SEAP and luciferase measurements. Statistical analyses were performed using one-way ANOVA followed by Tukey’s multiple comparisons test. Normality was assessed using the Shapiro–Wilk test prior to parametric analysis.* Indicates p<0.05 compared to Control.

### Screening of a gut-microbiome metabolite library for novel compounds which can stimulate interferon signaling

We wanted to identify a molecule that could increase IFN signaling without increasing NF-κB signaling in response to low-dose LPS (100pg/ml LPS) and thus rebalance the two pathways. Patients with AH are chronically experiencing elevated endotoxin levels, so we wanted to find a molecule that could rebalance these two pathways in the context of low-dose LPS (100pg/ml LPS). We utilized a library of 152 gut microbiome-derived metabolites (Cayman Chemical, Ann Arbor, Michigan, USA). This library includes microbial metabolites categorized as amino acid derivatives, aromatic compounds, hydroxy- and branched-chain fatty acids, sphingolipids, and nucleotide metabolites. We challenged THP1-Dual cells with low dose LPS (+/- 100pg/ml) on day 1 to simulate chronic endotoxemia. After 24 hours, we added 10uM of an individual microbial metabolite. On day 3, we measured SEAP and luciferase levels from the supernatant.

In control (without LPS challenge), most metabolites from the library produced subtle effects on both SEAP and luciferase levels (**Figure 3a**). Several compounds including naringin, urolithin A and prunin showed higher levels of luciferase compared to the other compounds, with no impact on SEAP secretion. This result suggests that these metabolites preferentially modulate the interferon signaling pathway rather than the NF-κB pathway. On the other hand, compounds such as gallic acid and 3-hydroxy lauric acid strongly increased SEAP levels, but not luciferase, so these molecules exacerbate pro-inflammatory signaling even without LPS stimulus. Both are aromatic compounds, which is consistent with previous reports that aromatic amino acid-derived metabolites can modulate innate immune signaling pathways[36, 37]. Muramyl dipeptide highly activated both pathways, which is consistent with other studies showing that it can signal through oligomerization domain 2 (NOD2) and engages both NF-κB and IFN pathways[38, 39].When cells were stimulated with low-dose LPS (100pg/ml LPS), the diversity of responses increased markedly (**Figure 3b**). Although many metabolites affected both NF-κB and IFN signaling, 10 compounds showed a disproportionate rise in IFN activity relative to NF-κB. These included N-Formyl-Met-Leu-Phe, palmitoleic acid, 2-hydroxy myristic acid, 17-methyl stearic acid, 7-methylxanthine, octanoic acid, naringin, urolithin A, and 4-HPAA (**Figure 3b**). These compounds preferentially upregulated the IFN pathway without simultaneously activating the NF-κB pathway. Such selective modulation may represent an ideal profile for restoring immune pathway balance in AH patients. On the other hand, compounds such as sphingosine (d18:1) showed very high SEAP expression compared to other metabolites, with limited effect on luciferase, suggesting a compounding effect with low-dose LPS (100pg/ml LPS). This is consistent with previous reports that pro-inflammatory role of sphingolipid metabolites in TLR4 signaling[40].

**Figure 3:**
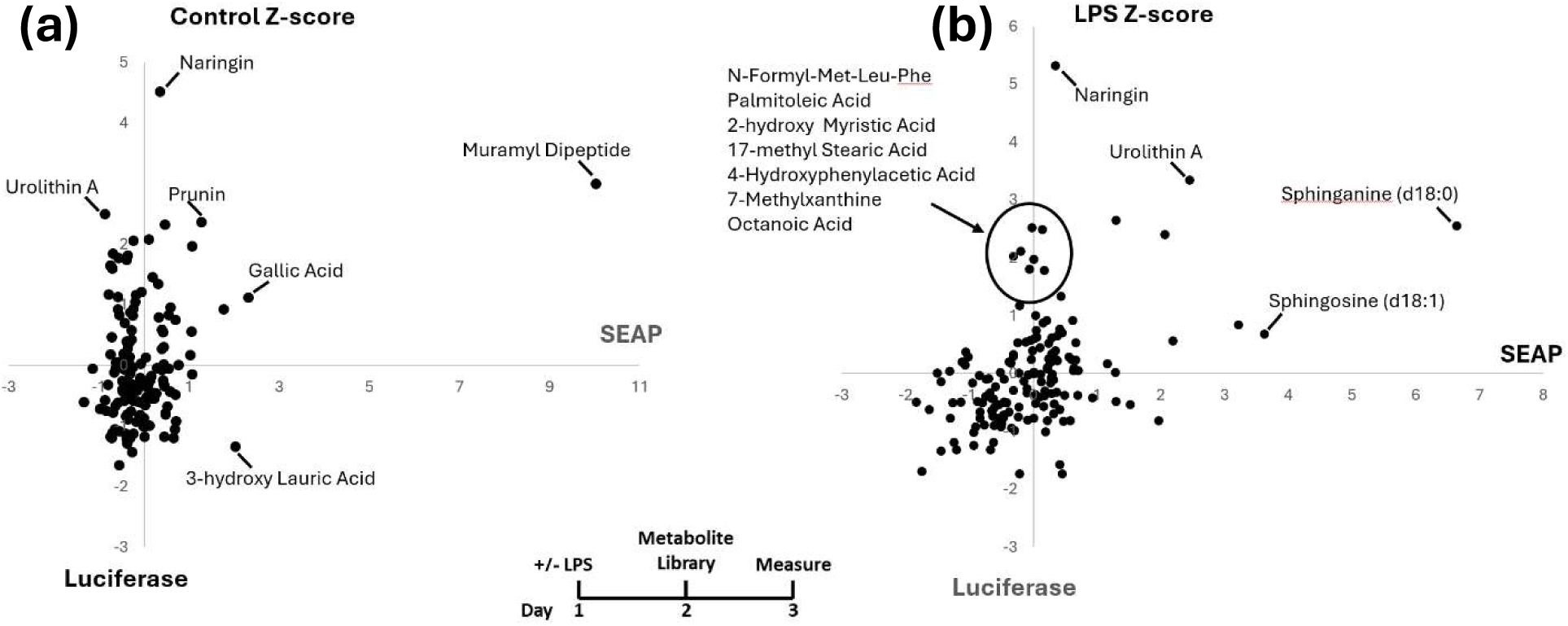
Z-score plot in both control and LPS group from metabolite library. (a) SEAP and luciferase results in Z-score in control group; (b) SEAP and luciferase results in Z-score in LPS group. Day 1, THP1-Dual cells were challenged with or without 100 pg/ml LPS; Day 2, THP1- Dual cells were treated with microbial metabolites; Day 3, cell supernatant was collected for SEAP and luciferase measurements.

Among all screened metabolites, 4-HPAA emerged as the most promising candidate. Among the 10 compounds that upregulated the interferon pathway after low-dose LPS (100pg/ml LPS), 9 of them had varying levels of luciferase activation in controls without LPS, suggesting baseline interferon activation (Figure S1, Table S1). Only 4-HPAA selectively enhanced IFN signaling, while exerting minimal NF-κB activation, in low-dose LPS (100pg/ml LPS) challenged cells and not controls (**Figure S1**). 4-HPAA is an aromatic amino acid–derived microbial metabolite produced during tyrosine fermentation[41], and has been found to inhibit weight gain[42]. Evidence has shown that 4-HPAA pretreatment protected against acetaminophen-induced acute liver injury in mice[43], and 4-HPAA markedly reverses hepatic fat accumulation in mice with steatosis[44]. Compared with other aromatic metabolites such as naringenin or urolithin A, which increased IFN expression under both control and LPS treatment, 4-HPAA uniquely favored IFN-dominant signaling only with LPS challenge. This pattern suggests that 4-HPAA may help restore balance between antiviral and inflammatory pathways by promoting IFN-driven responses while limiting NF-κB-mediated cytokine production.

After the screening test, we performed a dose response test of 4-HPAA to find an optimal working concentration. In vitro, 10 μM 4-HPAA was chosen as the most active concentration in our dose-response assay (**Figure S2**). However, this dose should be interpreted as a mechanistically active experimental concentration, since circulating 4-HPAA is typically detected at submicromolar levels in human serum, while local portal/hepatic exposure may be higher because of extensive first-pass metabolism[45].

### Dietary 4-HPAA modulates immune gene expression in a NIAAA-model

Next, we wanted to determine if 4-HPAA could reduce liver damage using a mouse model of ALD. We used the NIAAA chronic alcohol feeding model, where female mice are given a Lieber DiCarli diet with 5% ethanol for 10 days, then on the last day mice were given a binge of 5 g/kg ethanol by oral gavage[46]. Mice were sacrificed 6 hours after gavage, which is when peak inflammation, not liver damage, occurs[47]. Mice were divided into four groups: control, 4-HPAA, ethanol, and ethanol + 4-HPAA. For the mice given 4-HPAA, 0.4mg/ml 4-HPAA was added to the Lieber DiCarli diet. 4-HPAA was administered at 0.4 mg/mL (2.6 mM) in the Lieber-DeCarli diet. This dose was chosen to achieve sustained dietary exposure to a gut microbiota-derived metabolite and is within the range of concentrations previously used in mouse supplementation studies[48, 49]. We found that compared to ethanol group, ethanol + 4-HPAA group had more food intake after day 9 and more food intake in total (**Figure4b, Figure S3**). Ethanol feeding strongly elevated both ALT and AST compared to controls, confirming ethanol-induced liver injury (**Figure 4a, Figure S4, Table S2**). In ethanol-treated mice, ALT remained high regardless of 4-HPAA co-treatment, whereas AST levels in the ethanol + 4-HPAA group were reduced and less variable compared to ethanol alone. 4-HPAA alone did not elevate ALT or AST, indicating no liver toxicity under basal conditions. These findings suggest that while 4-HPAA did not normalize transaminase release, it may partially moderate ethanol-induced liver injury, particularly by lowering the magnitude and variability of AST responses.

**Figure 4:**
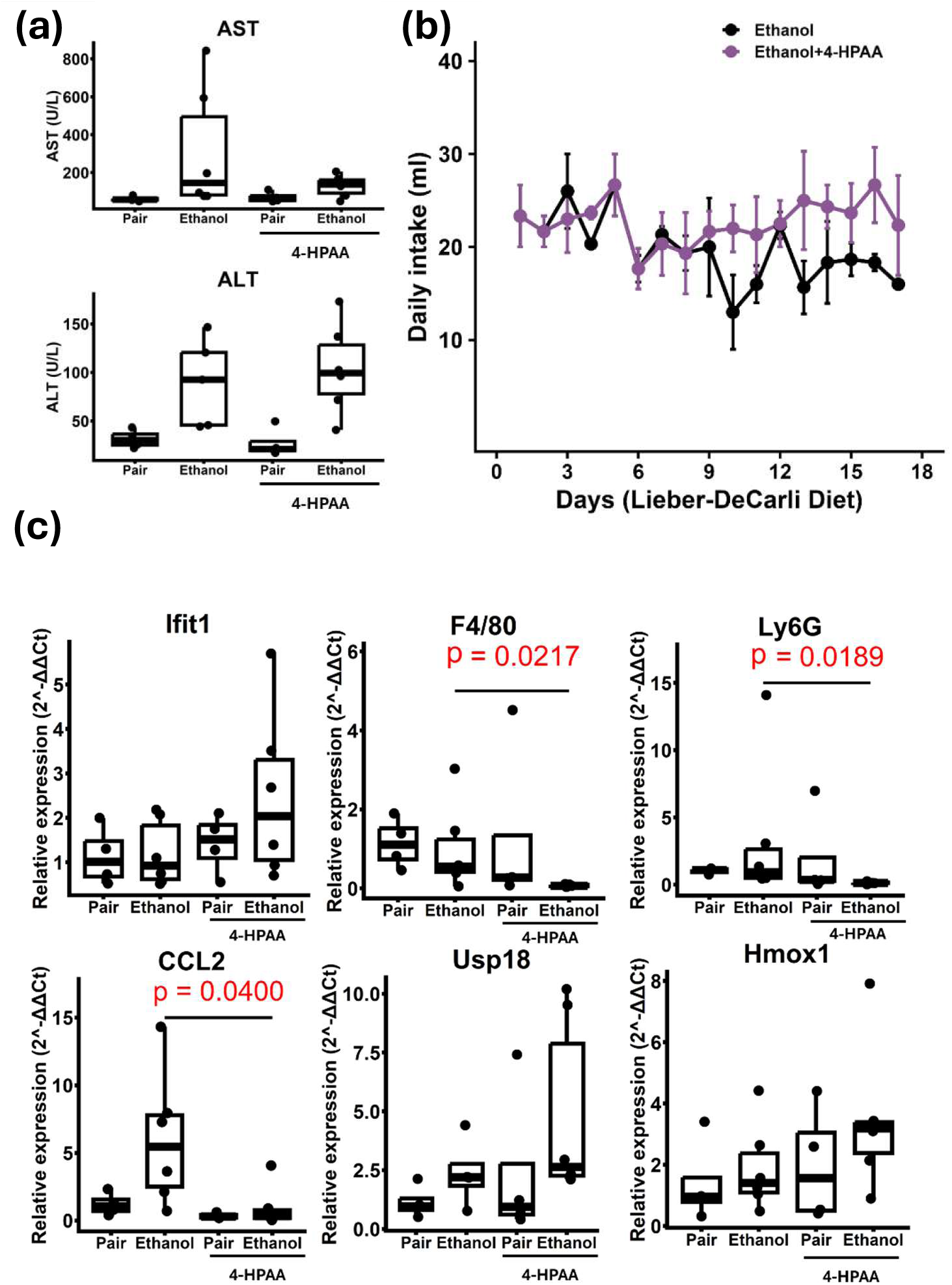
Effects of dietary 4-HPAA on liver injury and hepatic gene expression. (a) ALT and AST of plasma samples; (b) Food intake (ethanol vs ethanol + 4-HPAA); (c) qPCR results of dietary 4-HPAA test from mice liver samples. Statistical analyses were performed using one-way ANOVA followed by Tukey’s multiple comparisons test. Normality was assessed using the Shapiro–Wilk test prior to parametric analysis.

qPCR analysis of liver tissues (**Figure 4c**) revealed that co-administration of 4-HPAA with ethanol markedly increased *Ifit1* expression compared to the ethanol-only group. Markers of immune cell infiltration, *Ly6g* (an infiltrating neutrophil marker) and *F4/80* (a macrophage marker), were significantly reduced with 4-HPAA supplementation in ethanol-treated mice compared to ethanol group. Similarly, the ethanol group exhibited the highest expression of *CCL2*, a chemokine that promotes monocyte recruitment, whereas 4-HPAA supplementation significantly attenuated this effect. In contrast, expression of *USP18*, a type I interferon–responsive regulatory gene, was modestly increased by e thanol and was most strongly induced in the ethanol + 4-HPAA group, suggesting enhanced interferon-associated feedback regulation. We also assessed the antioxidant stress response gene *Hmox1* (heme oxygenase-1). While ethanol feedingslightly increased *Hmox1* relative to controls, expression was most strongly upregulated in the ethanol + 4-HPAA group, suggesting that 4-HPAA possibly enhances antioxidant defense pathways *in vivo* (**Figure 4b**). Together, these findings indicate that 4-HPAA not only moderates ethanol-induced liver injury but also reshapes immune and stress-response signaling toward a more protective profile.

### 4-HPAA reverses ethanol-driven shifts in cecal lipid metabolism

Given that 4-HPAA is produced by gut microbes, we wanted to know how 4-HPAA affects the gut metabolome and if it can reverse the damage caused by alcohol. We performed metabolomics analysis on mouse cecum samples to further explore the impact of ethanol feeding and 4-HPAA supplementation. A total of 6915 different metabolites were found from all samples. First, we explored the effect of alcohol on gut metabolome. A total of 153 metabolites were significantly changed between pair and ethanol groups (**Figure 5a**). Among them, 95 metabolites were significantly increased (*P* < 0.05) and 58 were significantly decreased (P<0.05) in the ethanol group relative to pair-fed controls. Among these metabolites, around 54% were unidentified. But among identified metabolites, top 3 were phosphatidylcholine (PC) lipids and were upregulated by ethanol (**Figure 5b**). Interestingly, many unidentified metabolites co-clustered with PCs and other lipid species, indicating that ethanol-induced changes extend beyond annotated compounds and broadly affect lipid families. In network figure, node size corresponds to VIP scores from the PLS-DA model, indicating the relative contribution of each feature to group separation. The clustering patterns therefore suggest that metabolomic differences are driven by coordinated shifts in molecular families, rather than changes in individual metabolites. Network analysis of the top 3 significant metabolites (**Figure 5e**) further demonstrated that these discriminating features are not independent, but instead cluster together within a PC–dominated lipid family. PC is one of the most abundant and essential phospholipids and a major component of biological membranes[50]. Many of the top metabolites were annotated as PC analogs with varying fatty acid side chains, suggesting that ethanol consumption drives coordinated remodeling of PC lipids. This clustering highlights that the ethanol effect is not restricted to isolated compounds but extends across a family of structurally related membrane lipids that share common fragmentation patterns.

**Figure 5:**
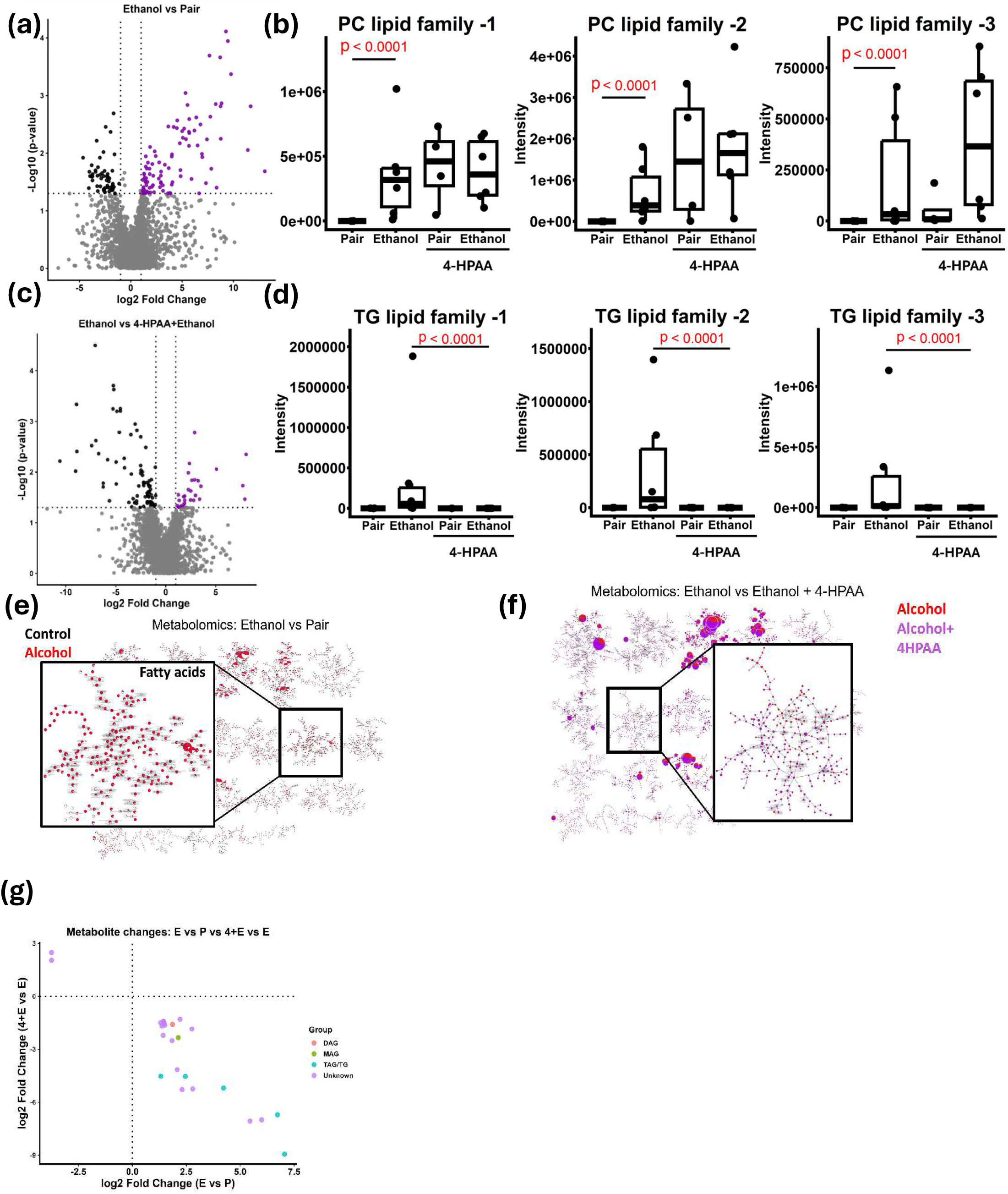
Metabolomics analysis on mouse cecum samples. (a) Volcano plot of metabolites (Ethanol vs Pair); (b) Box plots (Ethanol vs Pair); (c) Volcano plot of metabolites (Ethanol vs 4- HPAA + Ethanol); (d) Box plots (Ethanol vs 4-HPAA + Ethanol); (e); Molecular Community Network figure of comparison between ethanol and pair; (f) Molecular Community Network figure of comparison between ethanol and 4-HPAA + ethanol;(g) Common metabolite changes in both ethanol vs pair and ethanol vs 4-HPAA + ethanol. Statistical analyses were performed using one-way ANOVA followed by Tukey’s multiple comparisons test. Normality was assessed using the Shapiro– Wilk test prior to parametric analysis.

In contrast, 4-HPAA reversed many of the changes observed in ethanol groups. Differential metabolite analysis (**Figure 5c**) showed 80 significantly downregulated (*P* < 0.05) metabolites upon 4-HPAA supplementation when compared to ethanol alone group, partially reversing ethanol-induced elevations. 31 metabolites were significantly increased (*P* < 0.05) with 4-HPAA treatment in the ethanol group. Among these metabolites, around 72% of metabolites were unidentified. Top 3 known metabolites abundances across groups were shown in the **Figure 5d**. Network analysis of top 3 significant metabolites (**Figure 5f**) revealed that these features mapped predominantly to triacylglyceride (TG) clusters, a family of neutral lipids involved in energy storage[51]. Importantly, the clustering patterns indicate that these metabolites represent structurally related analogues rather than independent compounds, reflecting coordinated changes within a lipid family. The fact that multiple TG analogs were coordinately downregulated suggests that 4-HPAA exerts a targeted effect on lipid metabolism by counteracting ethanol-induced accumulation of triglycerides. Among the 80 metabolites altered by 4-HPAA, 24 were altered by EtOH, suggesting a reversal caused by 4-HPAA. Although most metabolites remained unidentified, 7 metabolites were identified under the glycerolipid family. Among these 7 metabolites, 5 metabolites were triacylglycerol (TAG), 1 metabolite was diacylglycerol (DAG), and 1 metabolite was monoacylglycerol (MAG) (**Figure 5g**). All these metabolites showed significant reduction in 4-HPAA treatment with ethanol compared to ethanol alone group. At the same time, all 7 metabolites were increased in mice when they had ethanol diet compared to pair-fed group. Previous reports also showed that ethanol consumption increased TAG/TG in mice liver[52]. These results suggested that 4-HPAA can reduce effects on glycerolipid accumulation which was brought by ethanol feeding.

We also compared pair with pair + 4-HPAA and pair + 4-HPAA with ethanol + 4-HPAA. Differential metabolite analysis showed that there were 350 and 392 significantly expressed metabolites respectively (**Figure S5**). Interestingly, in pair-fed mice, 4-HPAA still decreased multiple glycerolipids/TG-related metabolites, consistent with the alcohol-fed mice. Compared to pair-fed + 4-HPAA, ethanol also elevated PC abundance even with 4-HPAA supplementation.

In summary, these results point to a complementary pattern of lipid remodeling: ethanol feeding primarily elevates phosphatidylcholine lipids, while 4-HPAA supplementation selectively decreases TG. Both effects are evident not only at the level of individual metabolites but also across structurally related lipid families, underscoring that these metabolic changes are coordinated within chemical classes rather than being isolated molecular events. This family- level organization suggests that ethanol and 4-HPAA act on distinct but interlinked branches of hepatic lipid metabolism, with PCs reflecting ethanol-induced membrane remodeling and TGs reflecting 4-HPAA–mediated modulation of energy storage lipids.

### 4-HPAA modulates immune signaling in BMDM

To further explore 4-HPAA effects on other immune cell types, we tested 4-HPAA using bone marrow-derived macrophages (BMDMs) from mice to understand how it modulates innate immune signaling in primary macrophages. 8 mice were sacrificed for BMDM isolation (4 males and 4 females). After isolation of bone marrow cells, we differentiated cells for 10 days using 20ng/ml M-CSF to get BMDMs. On day 11, we treated cells with 4-HPAA (+/- 10uM). On day 12, we harvested cells and extracted RNA for RNA-seq analysis.

Differential gene analysis showed that there is a significant difference between control and 4-HPAA group (**Figure 6a-b, Table S3**). Transcriptomic analysis revealed that 4-HPAA treatment triggered a coordinated upregulation of extracellular matrix (ECM) related genes in macrophages (**Figure 6c**). Multiple fibrillar and basement membrane collagen genes, including *Col1a1, Col1a2, Col3a1,* were signifi cantly increased in 4-HPAA–treated cells compared with controls, indicating enhanced expression of ECM components[53]. In addition, genes involved in collagen post-translational modification and matrix stabilization were upregulated, including *Plod2*, which catalyzes collagen lysine hydroxylation required for fibril maturation[54], and *Lox* along with *Loxl1* and *Loxl2*, which mediate collagen cross-linking and contribute to matrix stiffening[55]. Consistent with active ECM turnover, 4-HPAA also increased expression of matrix metalloproteinases (*Mmp2* and *Mmp13*), enzymes that degrade collagen and facilitate matrix remodeling[56], together with their endogenous inhibitors *Timp1* and *Timp2*, which regulate proteolytic activity and balance matrix deposition versus degradation[57]. Collectively, these transcriptional changes define an ECM remodeling and tissue repair–associated macrophage program, suggesting that 4-HPAA modulates macrophage functions that influence the extracellular microenvironment. On the other hand, several immune activation and recruitment genes were significantly downregulated in the 4-HPAA group (**Figure 6c**), including *Clec4d*, *Ccr1*, *C3ar1*, *Tlr7*, and *Trem1*. These genes are related to innate immune sensing and inflammatory amplification (*Tlr7*and *Clec4d*), leukocyte chemotaxis (*Ccr1*), and complement-driven inflammatory signaling (*C3ar1*), while *Trem1*is a well-known amplifier of myeloid inflammatory responses. This result suggested that a less inflammatory response caused by 4-HPAA in macrophages. This result suggested that 4-HPAA induced less inflammatory response in macrophages.

**Figure 6:**
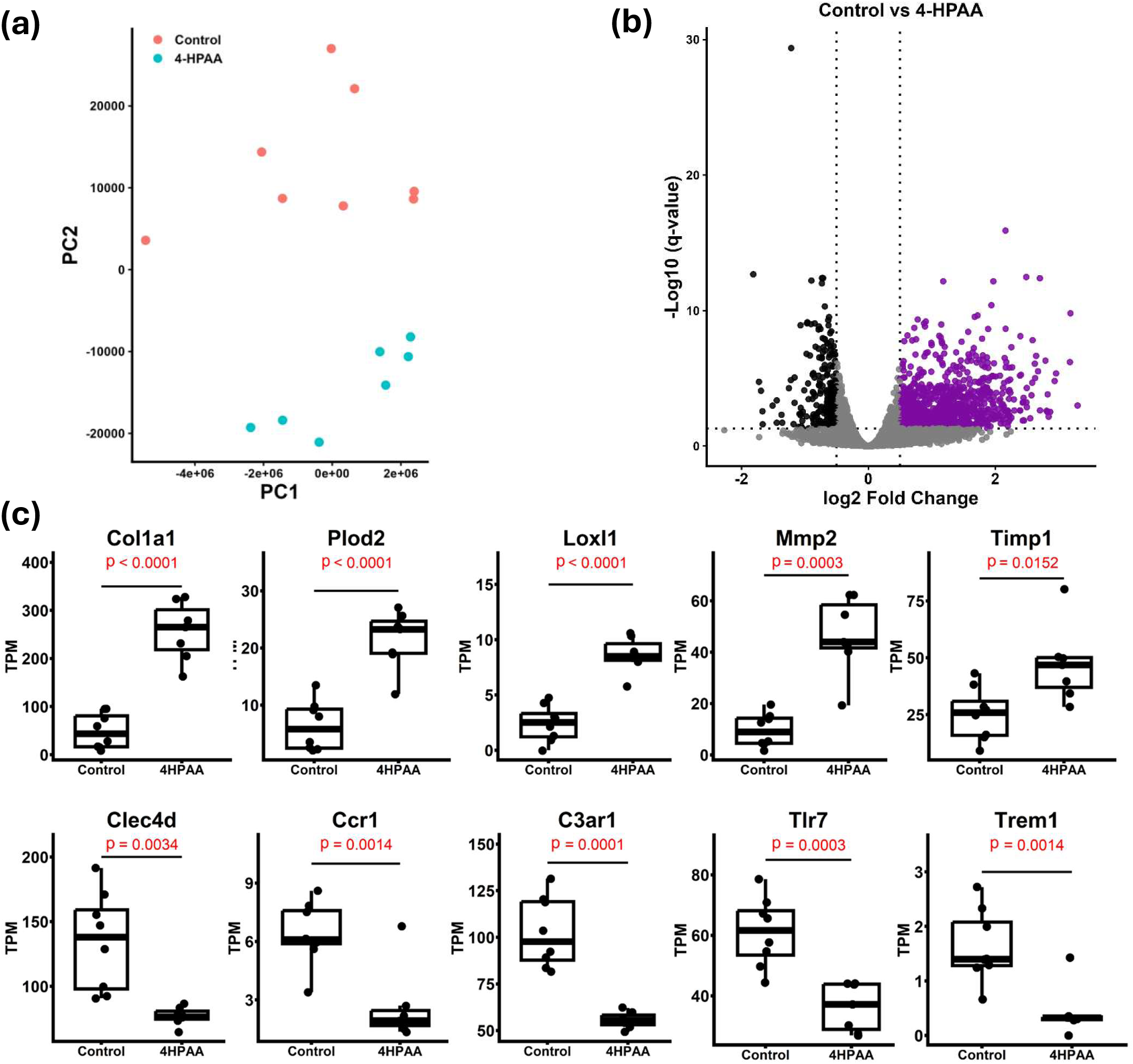
Comparison between control and 4-HPAA group. (a) PCA plot; (b) Volcano plot; (c) Box plots of specific genes. Statistical analyses were performed using unpaired two-tailed Student’s t-tests. Normality was assessed using the Shapiro–Wilk test prior to parametric analysis

Next, in order to define sex-dependent macrophage responses to 4-HPAA, differential expression and functional enrichment analyses were performed separately in female and male samples (**Table S4, Table S5**). As shown in **Figure 7a**, 4-HPAA treatment induced a coordinated upregulation of ECM related genes in macrophages, with stronger induction in females than in males. Female macrophages exhibited robust increases in multiple fibrillar and basement membrane collagen genes (*Col1a1, Col1a2, Col3a1, Col4a1/Col4a2,* and *Col6a1/Col6a2/Col6a3*), as well as genes involved in collagen maturation and matrix remodeling (*Lox, Loxl1, Loxl2, Plod2, Mmp2,* and *Mmp13*). In contrast, male macrophages showed a more attenuated transcriptional response, with reduced effect sizes across the same ECM gene set. Consistent with these gene-level differences, functional enrichment analysis showed that genes upregulated by 4-HPAA were strongly enriched for ECM and adhesion/cytoskeletal programs, including extracellular matrix organization, collagen formation, focal adhesion, and regulation of cell–substrate adhesion (**Figure 7b**). In contrast, downregulated genes were enriched for inflammatory and innate immune activation pathways, including leukocyte/neutrophil degranulation, lysosome/phagocytosis-related processes, MAPK signaling, cytokine/IL-6 production, and ROS/RNS production in phagocytes. Together, these results indicate that 4-HPAA promotes an ECM remodeling/tissue-repair transcriptional program while concomitantly dampening inflammatory effector responses in macrophages.

**Figure 7:**
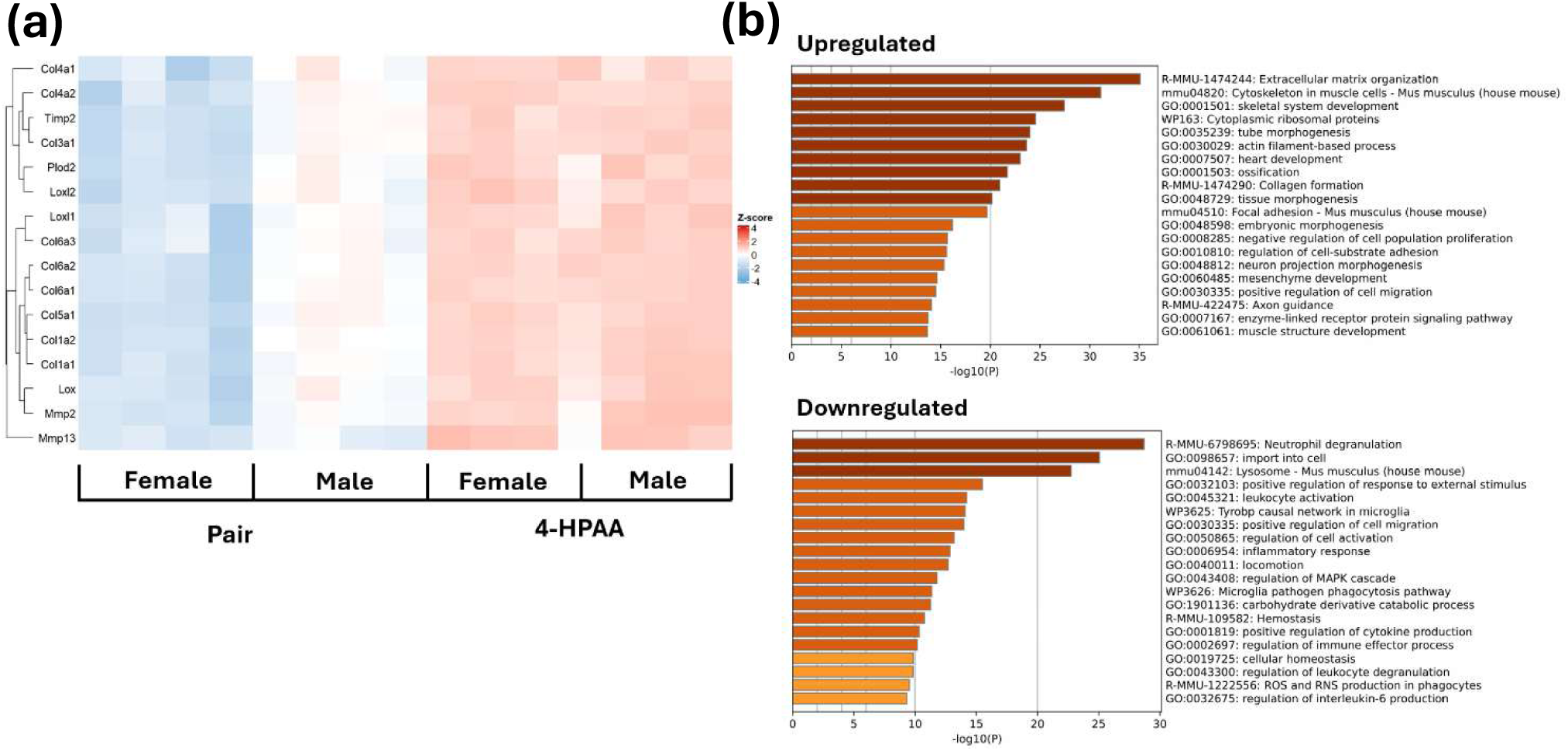
Comparison between control and 4- HPAA group by sex. (a) ECM/collagen pathway genes; (b) Enrichment analysis

### 4-HPAA enhances NRF2 signaling

To further explore these results, we reanalyzed the liver RNA-sequencing dataset from Osborn et al.[23]. They treated obese mice with 4-HPAA and performed bulk liver RNA-seq, which revealed transcriptional reprogramming characterized by reduced inflammatory and lipogenic gene expression and enrichment of fatty acid metabolism pathways consistent with reversal of hepatic steatosis. While their original analysis focused primarily on lipid metabolism and AMPK signaling, our secondary analysis revealed additional immune-related transcriptional changes. Data showed that 4-HPAA and control group have distinct differences (**Figure 8a-b**). In a transcriptomic reanalysis of liver samples, 45 genes were significantly altered by 4-HPAA treatment compared to controls (FDR < 0.05, **Table S6**). 4-HPAA upregulated several detoxifying/antioxidant genes related to the NRF2 pathway. For example, *Gstp1*, a canonical NRF2 target gene involved in glutathione-dependent detoxification[58], was significantly upregulated by 4-HPAA treatment, further supporting activation of NRF2-mediated antioxidant defense pathways. *Ugt2b37*, a downstream gene of NRF2 pathway encodes a phase II detoxification enzyme which functions downstream of NRF2-associated xenobiotic clearance pathways[59], was also upregulated compared to control. *Ce2b*, which hydrolyzes esterified xenobiotics and lipid-derived metabolites, was significantly upregulated compared to control samples (**Figure 8c**). Although *Ces2b* is not a canonical NRF2 target gene, its upregulation is functionally linked to *NRF2* activation, as CES family esterases participate in downstream detoxification programs that are frequently co-induced with NRF2-regulated antioxidant and phase II metabolism pathways[60]. In our alcohol model, *Hmox1* was also upregulated by 4-HPAA supplementation. Interestingly, *Ifi27l2a*, an interferon stimulated gene, was significantly downregulated under 4-HPAA treatment compared to control. This may suggest attenuation of interferon-associated inflammatory signaling[61] in the high-fat diet context. Taken together, these findings demonstrate that 4-HPAA activates genes of the *NRF2*-regulated antioxidant and detoxification network across both ethanol-induced and high-fat diet–associated liver injury models. Meanwhile, 4-HPAA may also create suppression effects of inflammatory interferon signaling in HFD-associated liver injury models.

**Figure 8:**
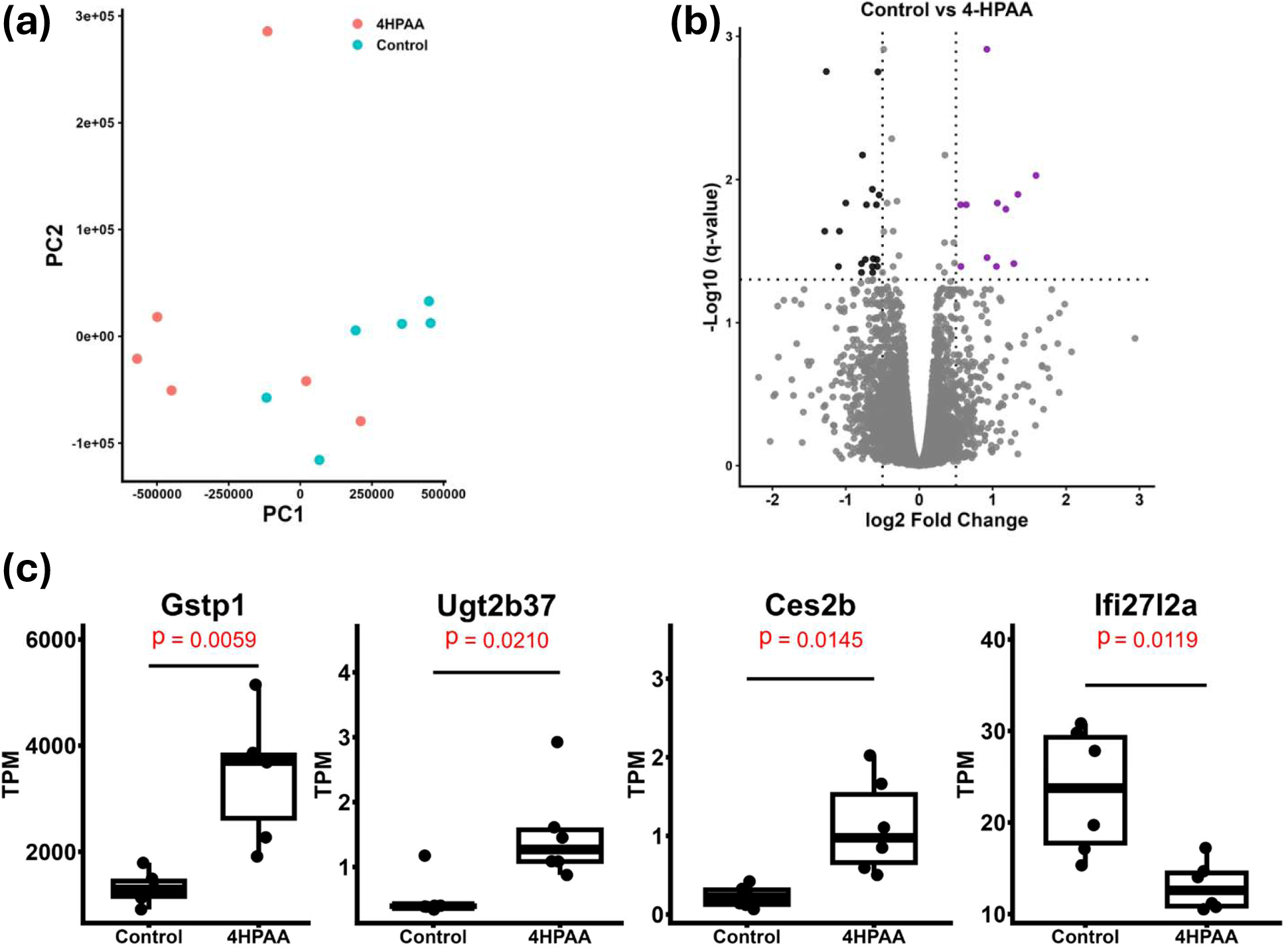
Reanalyzed RNA-seq data from Osborn et al. (n=6) (a) PCA plot by condition; (b) Volcano plot of differential expressed genes between control and 4-HPAA group; (c) Box plot of specific genes. Statistical analyses were performed using unpaired two-tailed Student’s t-tests. Normality was assessed using the Shapiro–Wilk test prior to parametric analysis

## Discussion

Our findings underscore a profound immune imbalance in alcohol-associated liver disease and demonstrate the potential of a microbiome-derived metabolite to correct it. First, analysis of liver transcripts from AH patients revealed an attenuated interferon-stimulated gene (ISG) response compared to other liver diseases, despite high inflammation. Notably, AH livers showed elevated expression of *SOCS1*, a potent feedback inhibitor of JAK-STAT signaling, alongside reduced *USP18*, which could reduce interferon signaling. This pattern suggests that chronic alcohol exposure creates a liver environment in which antiviral IFN pathways are suppressed. AH patients can become vulnerable to infections and limit tissue repair. Thus, alcohol’s effect on the immune system appears to shift the balance toward inflammation at the expense of antiviral defenses.

Using an *in vitro* model, we confirmed that short term ethanol exposure exacerbates proinflammatory signaling while suppressing interferon responses when cells encounter endotoxin. In THP1-Dual monocytes, even low-dose LPS (100pg/ml LPS) triggered NF-κB and IFN pathways; however, ethanol pre-treatment caused a dose-dependent increase in NF-κB activity (SEAP) and a decrease in IFN activity (luciferase reporter). Although ethanol was evaluated up to 100 mM to define dose-dependent effects, the physiological relevance is greatest at lower concentrations such as 10–20 mM, while the higher doses should be viewed as exploratory mechanistic conditions rather than direct equivalents of moderate alcohol consumption. These data indicate that ethanol synergizes with endotoxin to amplify TLR4/MyD88–NF-κB signaling, potentially through priming of inflammatory pathways or oxidative stress, as suggested by prior studies[62]. At the same time, ethanol impairs interferon production[63]. Biologically, this dual effect of ethanol provides a basis for immune dysregulation in AH.

To identify microbiome-derived metabolites with immunomodulatory activity, by screening a library of microbiome-derived metabolites, we identified several candidates that influenced NF-κB and IFN pathways. 4-HPAA emerged as the most promising hit from this screen. Uniquely, 4-HPAA selectively boosted interferon signaling with minimal effect on baseline or LPS-induced NF-κB activity. This suggests that 4-HPAA acts only under inflammatory stimulation, a desirable feature for a therapy aimed at pathological inflammation without over-activating the immune system in resting conditions. Moreover, 4-HPAA is an aromatic microbial metabolite derived from the gut metabolism of dietary polyphenols such as kaempferol and related flavonoids[64], linking diet, microbiota, and host immune regulation. Some of these other dietary polyphenols, like kaempferol and naringenin were also positive hits in our screen like 4-HPAA, suggesting a common role for polyphenols in protecting the liver. Previous studies reported that 4-HPAA can protect against acetaminophen-induced acute liver injury[43] and reverse hepatic steatosis in obese mice[42]. These known hepatoprotective properties complement our findings and raise the intriguing possibility that 4-HPAA’s benefits in ALD may be multisided, affecting both immune and metabolic pathways. Our screening strategy identified metabolites based on their functional effects on immune signaling pathways. However, we did not directly assess whether these metabolites are differentially abundant under alcohol-associated conditions. Because gut microbiome–derived metabolites exist within a complex and dynamic mixture in vivo, their biological effects are likely influenced by concentration, tissue exposure, and interactions with other metabolites. Therefore, the extent to which the observed effects of individual metabolites, including 4-HPAA, occur under physiological conditions remains to be determined.

In an in vivo model of alcoholic liver injury, 4-HPAA was added to the Lieber DiCarli diet containing ethanol. 4-HPAA, unlike many other flavonoids, is not thought to cross the gut barrier[23], but because alcohol consumption causes gut barrier disruption[65], we hypothesized that 4-HPAA in the diet should easily become systemically bioavailable in the alcohol-fed mice. Interestingly, 4-HPAA supplementation actually increased consumption of the alcohol-diet. This suggests that 4-HPAA improved tolerance to alcohol and any protective effects we observe are not due to less alcohol consumption or metabolism. 4-HPAA supplementation shifted hepatic immune responses toward an antiviral and anti-inflammatory state. Notably, 4-HPAA restored interferon pathway activity (increased *Ifit1* expression) despite ongoing ethanol exposure, suggesting it counteracts ethanol-induced immunosuppression and strengthens antiviral defenses. Simultaneously, 4-HPAA attenuated ethanol-driven inflammatory signals,notably downregulating key pro-inflammatory mediators (such as *CCL2*) as well as markers of neutrophil and macrophage infiltration (*Ly6G* and *F4/80*). This implies that 4-HPAA limits inflammatory leukocyte recruitment and activation, potentially reducing alcohol-induced tissue damage. 4-HPAA also upregulated the antioxidant enzyme *Hmox1* [66], indicating activation of cytoprotective stress responses (possibly via *NRF2*). Collectively, these findings reveal that 4-HPAA fosters a more protective immunological milieu in the alcohol-injured liver, highlighting a novel immunomodulatory mechanism in alcoholic liver disease.

Despite these favorable immunological effects, 4-HPAA produced only modest improvements in biochemical liver injury markers. In the ethanol-fed mice, 4-HPAA did not reduce ALT and only partially lowered AST, which showed a downward trend and less variability compared with ethanol alone. These findings suggest that 4-HPAA attenuates, but does not fully prevent, hepatocellular injury in this acute-on-chronic model given the severity of ethanol binge–induced damage. Importantly, 4-HPAA alone did not increase ALT or AST, indicating that it is not hepatotoxic at the administered dose. Although longer treatment or combination strategies may be required to achieve stronger protection, the partial improvement in AST along with reduced inflammatory and interferon-related dysregulation supports a hepatoprotective role for 4-HPAA.

While prior studies of alcoholic liver disease have focused primarily on hepatic or serum metabolites[67, 68], our analysis reveals that ethanol induces coordinated, family-level alterations in gut-derived lipid species. Metabolomic analysis supported the protective role of 4-HPAA by revealing that it corrects ethanol-induced lipid dysregulation. Chronic alcohol intake changed cecal lipid metabolism, exemplified by elevated PCs indicative of membrane remodeling under ethanol stress. PCs are major membrane phospholipids, and their increase suggests ethanol induces changes in gut or liver membrane composition, possibly through altered bile secretion[50] or microbial metabolism of lipids[69]. 4-HPAA largely reversed these changes, notably reduced the accumulation of triglyceride species associated with steatosis. This ability to reduce ethanol-elevated glycerolipids suggests that 4-HPAA counteracts a key metabolic feature of alcoholic liver disease (hepatic fat accumulation). Mechanistically, this metabolic shift might result from 4-HPAA enhancing lipid catabolism (AMPK activation) or altering gut microbial metabolism to decrease lipogenic substrate availability. Notably, 4-HPAA’s metabolic benefits coincided with increased hepatic *Hmox1* expression, hinting at reduced oxidative stress due to a lower lipid burden. The coordinated decrease in TGs and increase in antioxidant gene *Hmox1* with 4-HPAA might point to reduced oxidative stress and lipid peroxidation in the liver, since excessive triglyceride storage and lipotoxicity are known to generate oxidative injury in ALD[70, 71]. This complementary pattern underscores that 4-HPAA not only modulates immune pathways but also corrects metabolic imbalances, potentially tackling both inflammation and steatosis, two central features of alcoholic liver disease.

To examine the effects of 4-HPAA on other immune cell types, we treated BMDMs with or without 4-HPAA in the presence or absence of LPS. 4-HPAA treatment induced a clear shift in the BMDM transcriptome toward an ECM remodeling and tissue-repair–associated program. Although fibroblasts are classically considered the primary source of collagen deposition, accumulating evidence indicates that injury-activated macrophages can upregulate collagen genes and contribute directly to scar-associated ECM formation, consistent with a reparative macrophage state that supports wound healing. In parallel with ECM induction, 4-HPAA suppressed innate immune sensing and inflammatory amplification pathways, including reduced expression of *Clec4d*, *Ccr1*, *C3ar1*, *Tlr7*, and *Trem1*. This result suggests diminished capacity for inflammatory signal, chemotactic recruitment, and complement-driven activation. These gene-level patterns were consistent with pathway enrichment results indicating attenuation of pro-inflammatory processes in macrophages. Interestingly, female-derived macrophages exhibited stronger transcriptional responses than males across the same ECM and immune gene sets, highlighting sex as an important modifier of 4-HPAA responsiveness and suggesting potential contributions from hormone-dependent mechanisms.

Combining the findings from Osborn et al. and our alcohol model, these results highlight a consistent modulation of the NRF2 pathway by 4-HPAA across different models of liver injury. In NIAAA-model mice, 4-HPAA was previously shown to elevate *Hmox1* expression by qPCR, reflecting NRF2 pathway activation. Similarly, in the high-fat diet model reanalysis, 4-HPAA treatment induced NRF2-dependent cytoprotective genes such as *Gstp1*, indicating that 4-HPAA triggers a hepatic antioxidant response in both alcohol- and diet-induced steatosis contexts. This convergence suggests that 4-HPAA’s hepatoprotective effects are at least partly mediated by NRF2 activation and subsequent upregulation of antioxidant enzymes. The enhancement of such NRF2-driven defenses by 4-HPAA is consistent with reports that 4-HPAA can increase nuclear NRF2 and phase II/antioxidant enzyme activity *in vivo[43]*. Thus, both the chronic alcohol and high-fat diet models support a unifying mechanism whereby 4-HPAA enhances the liver’s antioxidant capacity via the NRF2 related pathway, which may contribute to its anti-steatotic and anti-inflammatory benefits.

In conclusion, this study provides evidence that 4-HPAA, a gut microbial metabolite, emerged as a potent immunometabolic regulator capable of counteracting ethanol’s pathogenic effects by rebalancing innate immune signaling and reducing lipid accumulation. These findings broaden our understanding of the interactive roles of microbial metabolites in liver disease and suggest that leveraging such metabolites could be a novel therapeutic avenue in alcoholic hepatitis. Direct supplementation would provide a more controlled and dose-defined approach, but future studies would be needed to establish safety, pharmacokinetics, tissue distribution, and optimal dosing. In addition to direct supplementation, an alternative strategy may involve modulating the gut microbiome to enhance endogenous 4-HPAA production, for example through targeted dietary interventions, prebiotics, or microbial consortia capable of producing 4-HPAA. Although microbiota-targeted strategies to increase 4-HPAA may be feasible, the specific microbial taxa and pathways responsible for endogenous 4-HPAA production are not yet fully defined and likely involve multiple substrate-dependent routes, including aromatic amino acid and flavonoid catabolism. With further validation and mechanistic elucidation, both 4-HPAA and microbiome-based approaches to increase its production could represent promising strategies to restore immune homeostasis and protect against alcohol-associated liver injury.

## Author Contributions

Y.Z., N.H., D.M., C.V., F.P., A.A. and A.K. contributed to the conception and design of the study. Y.Z., N.H., D.M., C.V., F.P., A.A and A.K. designed and performed all experiments. D.M., Y.Z., A.A. and A.K. analyzed metabolomics data. Y.Z. and A.K. analyzed RNA-seq data. Y.Z. wrote the firs draft of the manuscript. All authors contributed to manuscript revision, read and approved the submitted version.

## Acknowledgements

The authors thank the Clinical Research Unit, Genomics Core, and Computing Services at the Cleveland Clinic. This work was funded by the following: NIH-NIAAA grant: K99/R00AA028048 (A.K.).

## Data Availability

The bulk RNA-seq data from mouse bone marrow derived macrophage generated in this study can be found at National Center for Biotechnology Information Gene Expression Omnibus under accession number [TBD].

The bulk RNA-seq data from patient livers in this study can be found at dbGAP (phs001807.v1.p1)[20].

The bulk RNA-seq data from mouse livers can be found at National Center for Biotechnology Information Gene Expression Omnibus under accession number (GSE188967)[23].

## Code Availability

All scripts used for analyses can be found at author’s github (github/atomadam2).

