## Supplemental figures for "A microbial metabolite reduces alcohol-induced inflammation via dual modulation of NF-κB and Interferon pathway"

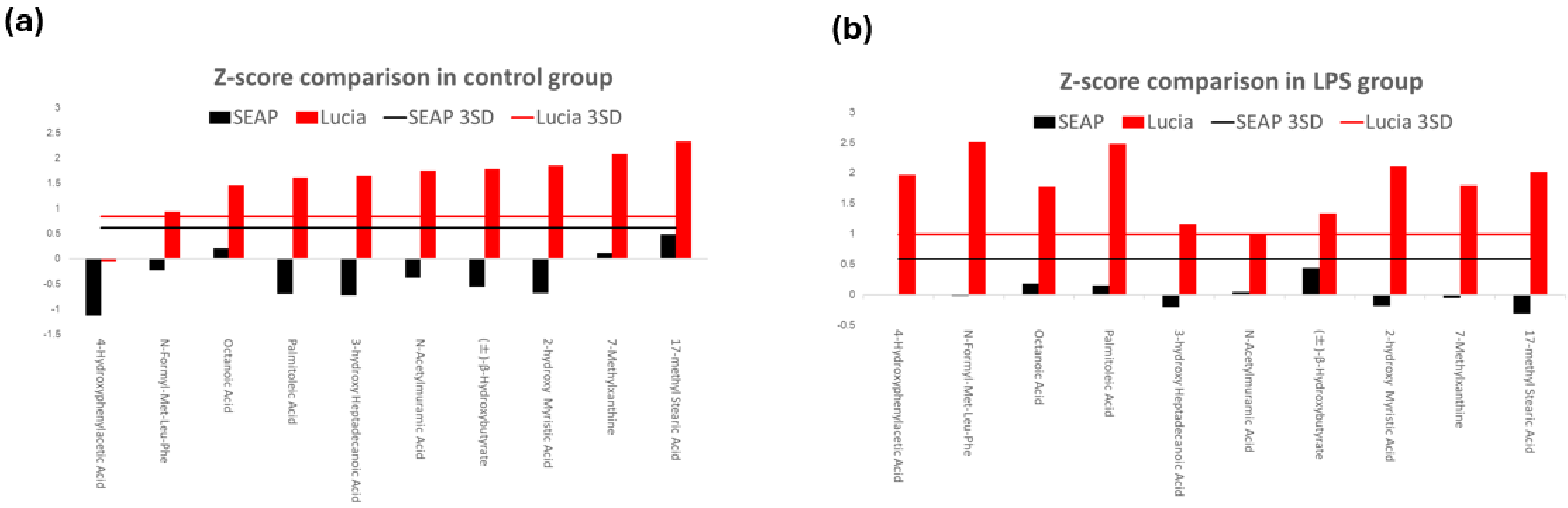


**Figure S1:** Z-score plot of 10 potential compounds. (a) Control group; (b) LPS group.

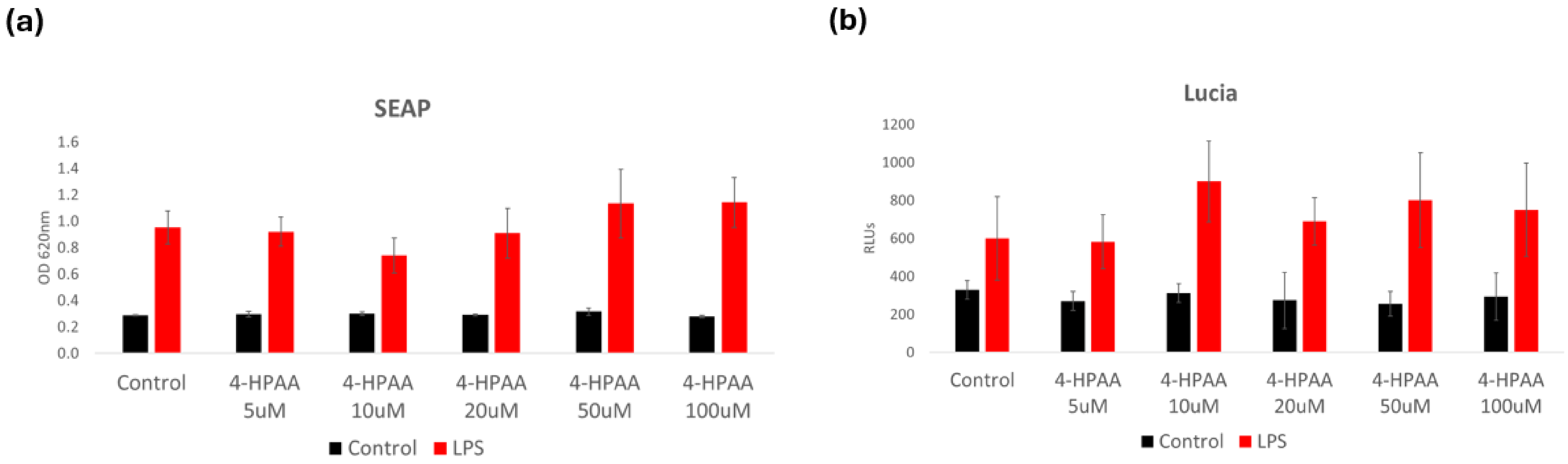


**Figure S2:** Does response test of 4-HPAA. (a) SEAP; (b) Luciferase.

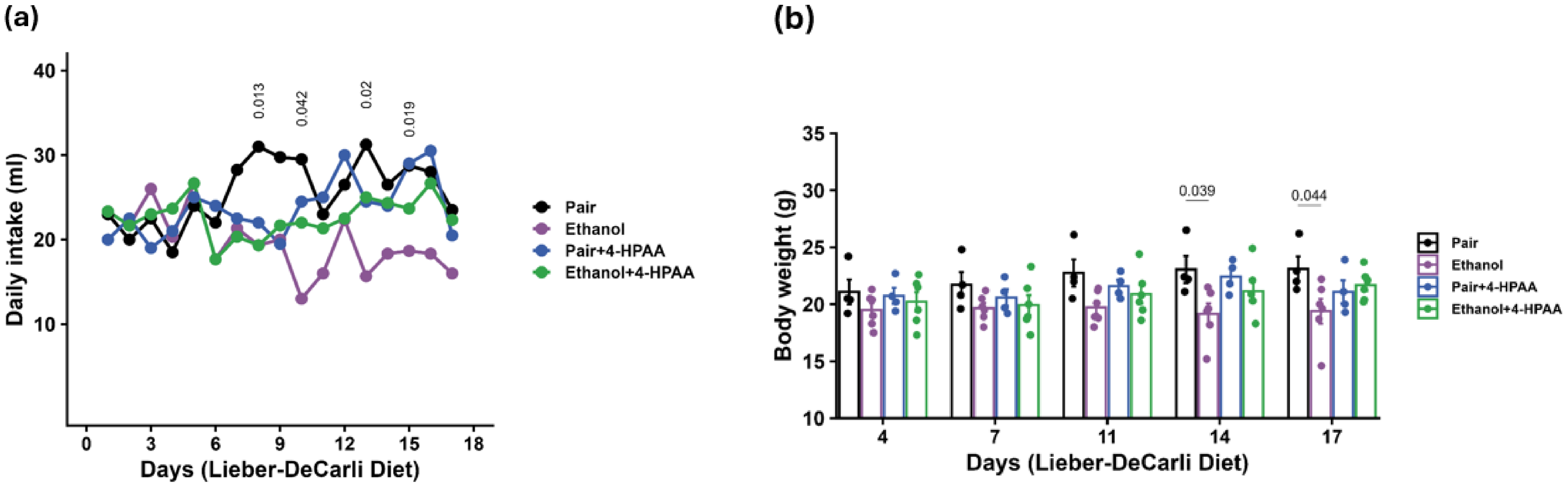


**Figures S3:** Effects of dietary 4-HPAA in mice. (a) Food intake; (b) Body weight

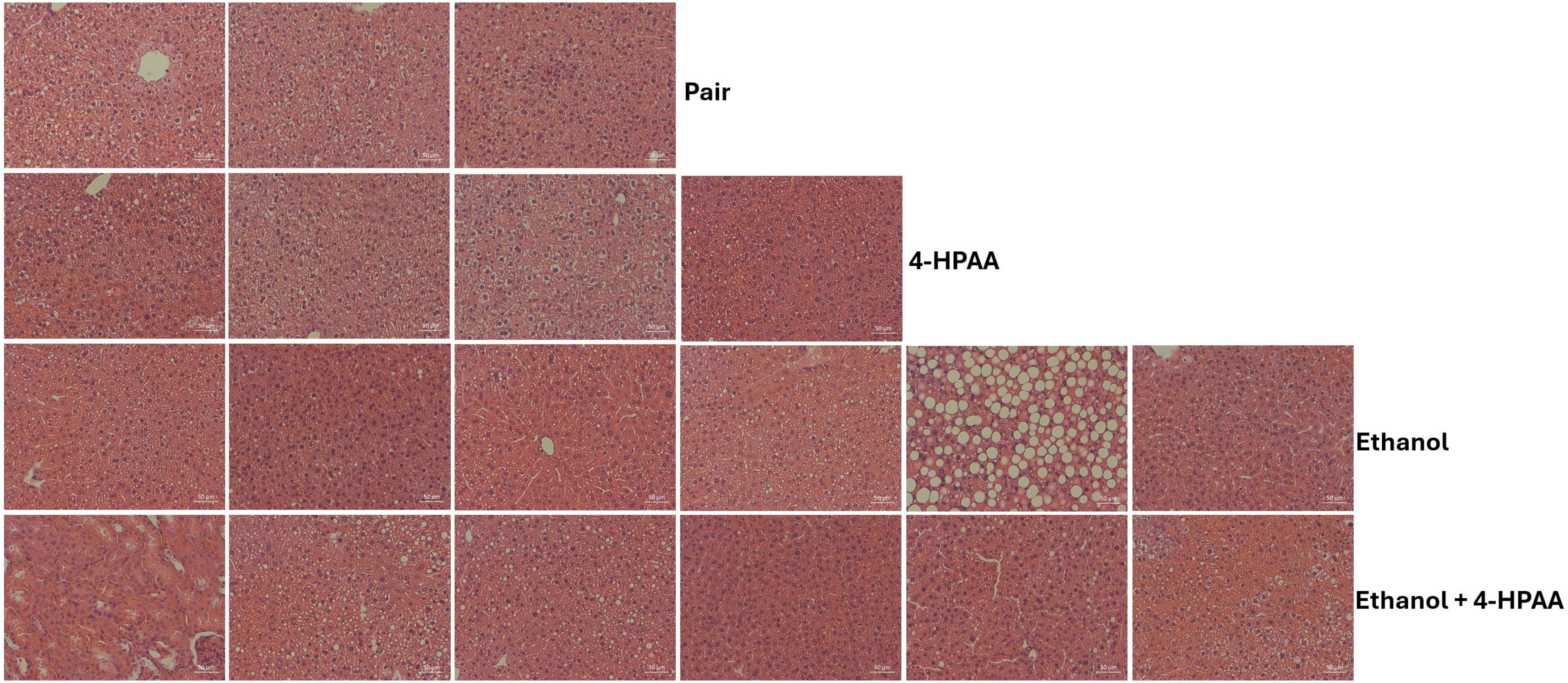


**Figure S4:** Representative H&E-stained liver sections from mice in the NIAAA chronic ethanol model.

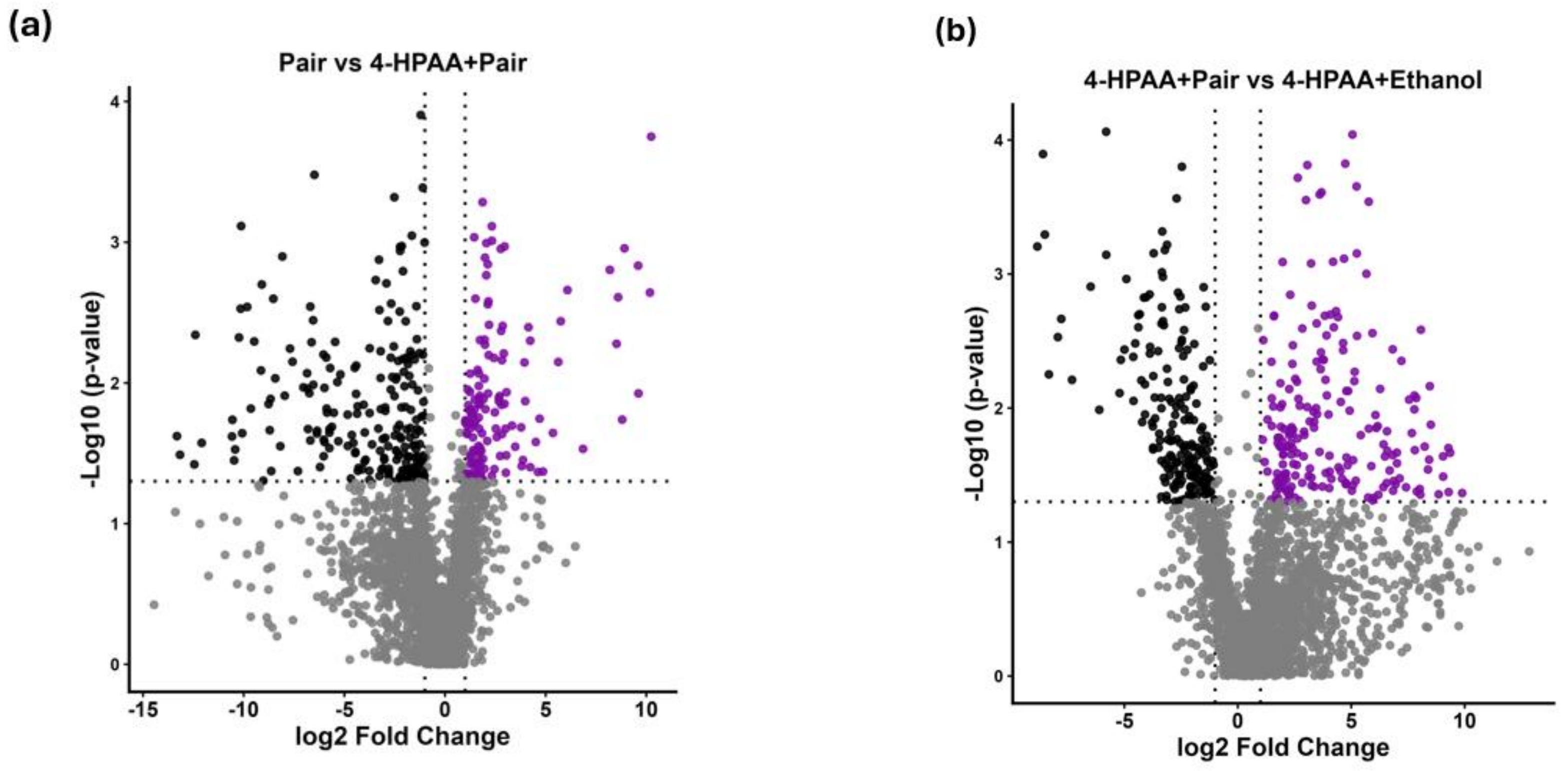


**Figure S5:** Differential gene volcano plot of dietary 4-HPAA mice experiment. (a) Pair vs 4-HPAA; (b) 4-HPAA vs 4-HPAA with ethanol.
